# Brain transcriptional regulatory architecture and schizophrenia etiology converge between East Asian and European ancestral populations

**DOI:** 10.1101/2021.02.04.922880

**Authors:** Sihan Liu, Yu Chen, Feiran Wang, Yi Jiang, Fangyuan Duan, Yan Xia, Zhilin Ning, Miao Li, Wenying Qiu, Chao Ma, Xiao-Xin Yan, Aimin Bao, Jiapei Dai, Richard F. Kopp, Liz Kuney, Jufang Huang, Shuhua Xu, Beisha Tang, Chunyu Liu, Chao Chen

## Abstract

Understanding the genetic architecture of gene expression and splicing in human brain is critical to unlocking the mechanisms of complex neuropsychiatric disorders like schizophrenia (SCZ). Large-scale brain transcriptomic studies are based primarily on populations of European (EUR) ancestry. The uniformity of mono-racial resources may limit important insights into the disease etiology. Here, we characterized brain transcriptional regulatory architecture of East Asians (EAS; n=151), identifying 3,278 expression quantitative trait loci (eQTL) and 4,726 spliceQTL (sQTL). Comparing these to PsychENCODE/BrainGVEX confirmed our hypothesis that the transcriptional regulatory architecture in EAS and EUR brains align. Furthermore, distinctive allelic frequency and linkage disequilibrium impede QTL translation and gene-expression prediction accuracy. Integration of eQTL/sQTL with genome-wide association studies reveals common and novel SCZ risk genes. Pathway-based analyses showing shared SCZ biology point to synaptic and GTPase dysfunction as a prospective pathogenesis. This study elucidates the transcriptional landscape of the EAS brain and emphasizes an essential convergence between EAS and EUR populations.

## Main

Population genetics examines differences within and between populations and how such genetic differences contribute to health and disease. A global understanding of the influence of genetic variance on complex diseases would advance insight into the biological mechanisms of disease risk^1^. During the past decade, genome-wide association studies (GWAS) have identified thousands of risk variants for psychiatric disorders across diverse populations^2,3^. Nonetheless, most of samples in psychiatric disorders GWAS originate from those of European (EUR) descent^4^. Due to ancestral differences evident in allele frequencies (AF), linkage disequilibrium (LD) patterns, and other factors, GWAS findings often fail to translate to other populations^5,6^. For example, Martin et al. examined that genetic risk prediction accuracy will decrease within heterogeneous populations which the original GWAS sample and target of prediction are divergent^5^.

Interpreting GWAS “hits” with expression quantitative trait loci (eQTL) and splicing quantitative trait loci (sQTL), significantly enriched for trait-associated SNPs offers a feasible alternative for advancing our understanding of the molecular mechanisms underlying complex traits^7,8^. In the past decade, eQTL and sQTL have become familiar and effective tools enabling GWAS to explain single nucleotide polymorphism (SNP) heritability and spotlighting potential disease risk genes^9–14^. Various methods have been proposed to interpret GWAS using eQTL/sQTL signals to establish gene-expression prediction models. Examples such as PrediXcan^15^ and TWAS^16^ correlate imputed gene expression to a phenotype under investigation. However with eQTL/sQTL, the problem of population disparity becomes even more extreme, as most resources focus largely on the EUR ancestry alone^17–20^. The capacity of existing prediction models to isolate causal genes common across populations appears to be constrained by the Eurocentricity of the models themselves. In this way, the limited availability of non-EUR GWAS impedes our ability to fully understand the genetic basis of diseases and to translate basic research into clinical medicine.

Recent studies have discovered significant inefficiency in predictive performance between heterogenous populations^14,21–23^. One plausible explanation for this is the differences in LD patterns and AF distribution. These disparities also hinder the ability of QTL to replicate in diverse populations. For example, comparing the regulatory architecture of gene expression in lymphoblastoid cell lines, Stranger *et al.* found that QTL differentiation among populations was likely due to AF differences reducing the statistical power of association testing^24^. Additionally, Lauren *et al.* showed that the AF differences between populations led to the accurate prediction of some genes and poor prediction in others^22^. Therefore, developing new transcriptome regulatory profiles and prediction models specific to ancestral populations is critical for accurately predicting gene expression and identifying disease risk genes.

It should be noted that transcriptomic studies conducted for other tissues (e.g., blood), cannot adequately represent the transcriptome of neuropsychiatric disorders that are most closely associated with the brain^25^. Gene expression is tissue-specific. Moreover, many studies have discovered that QTLs within specific pathogenic tissues are significantly enriched for relevant trait associations^26–30^. For example, in the frontal cortex, a region widely accepted as critical for schizophrenia (SCZ), QTLs detected are significantly enriched with greater SCZ GWAS signals than QTLs detected from other tissues^26–28^. Such findings signal the need to develop regulatory profiling of the human brain to uncover the biological mechanisms of SCZ. Several studies have generated large-scale postmortem brain data^31–33^. For instance, Wang *et al.* developed a comprehensive resource for functional genomics of 1,866 adult brains using PsychENCODE data that highlights key genes and pathways associated with SCZ, including immunological, synaptic, and metabolic pathways^33^. To our knowledge, no systematic investigation into whether the genetic control of gene expression and splicing in brain is similar or varies between populations exists. Furthermore, determining whether differences represent etiologic heterogeneity in SCZ across populations also begs investigation.

Here, we developed a novel brain transcriptome dataset comprised of 151 EAS individuals. We hypothesize that the brain’s regulatory architecture of gene expression and the etiology of SCZ converge between EAS and EUR populations. We also posit that AF and LD patterns distinct to ancestral populations are at least in part responsible for QTL heterogeneity across populations. To test these hypotheses, we applied eQTL, sQTL and co-expression analyses, comparing the results with existing PsychENCODE/BrainGVEX data (specifically the EUR subpopulation) to evaluate the similarities and differences in the transcriptional regulatory architecture of the two populations. By integrating eQTL and sQTL results with SCZ GWAS summary data, we quantified the enrichment of eQTL/sQTL associated with GWAS signals. Further, we identified common and novel risk genes as well as disease-related pathways. From these data we assembled a new genome-wide human brain regulatory map, which affords considerable insight into the biological progression of SCZ in East Asians.

## Results

To identify EAS-specific regulatory variants shaping brain gene expression and alternative splicing, we performed high-density genotyping and high-throughput RNA-sequencing in 151 EAS prefrontal cortices (Fig. 1). After quality control and preprocessing (Methods and Extended Data Fig. 1), we gathered 18,939 brain-expressed genes and 6.4 million autosomal SNPs. PCA (principal component analysis) of ancestry verified the East Asian ethnicity of all donors (Supplementary Note). eQTL and sQTL mapping and constructed co-regulatory networks enabled us to examine the brain expression regulatory architecture of each population individually, with the EUR population derived from the PsychENCODE/BrainGVEX project.

**Fig. 1.**
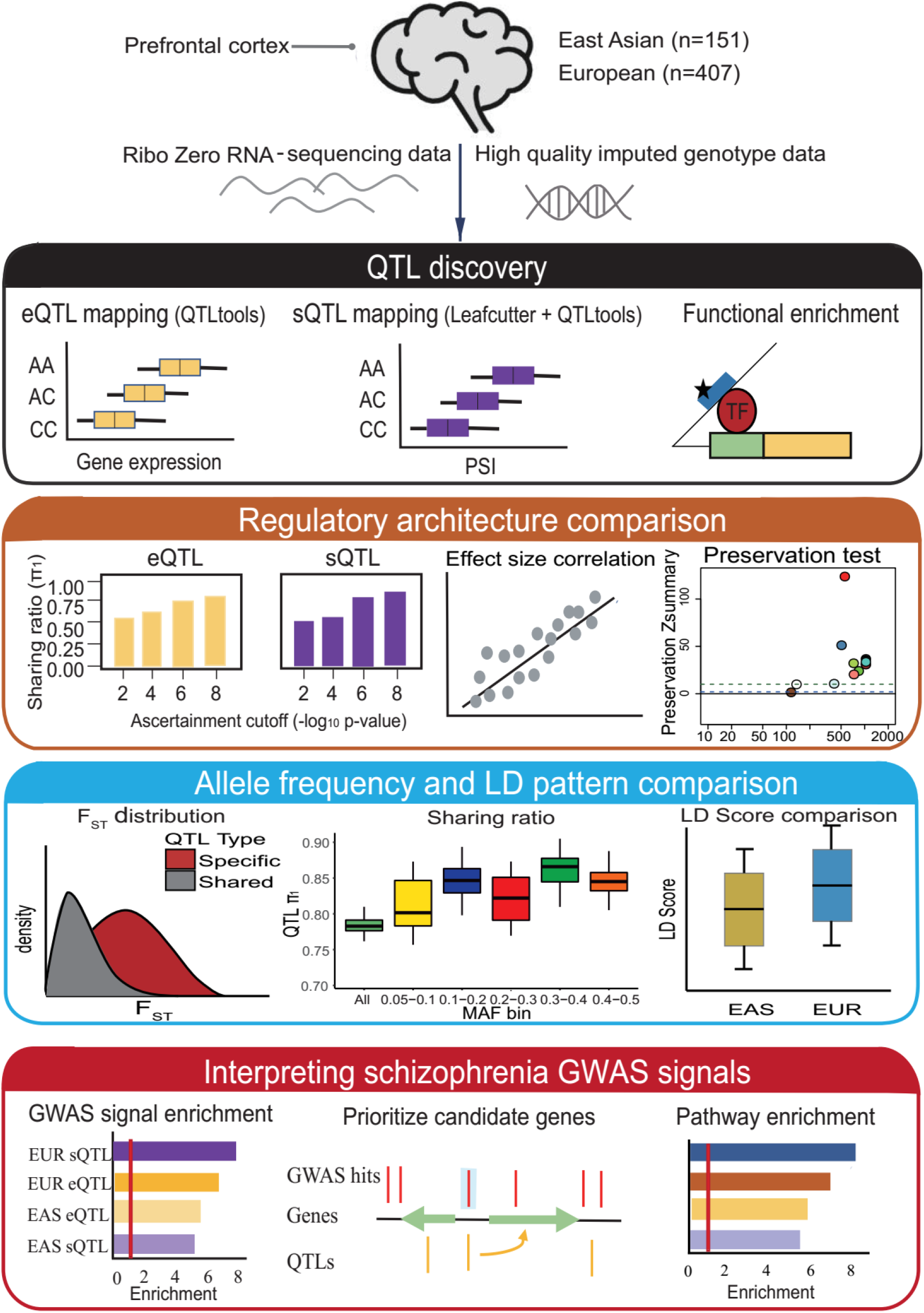
Study design. We collected genotype and RNA-seq data from East Asian (n = 151) and European populations (n = 407). After quality control and data preprocessing, eQTL and sQTL were independently calculated using standard methods of covariate correction. QTLs were characterized based on functional enrichment. Then, we compared the regulatory architecture including eQTL/sQTL and the gene co-regulatory patterns between EAS and EUR populations. Next, we calculated F_ST_ and LD scores to evaluate the contribution of AF and LD patterns difference in QTL comparison. Finally, to determine whether schizophrenia biology between East Asian and European populations is analogous, we integrated QTLs previous identified with SCZ GWAS to identify disease risk and important biological processes under genetic control. PSI: percent-spliced-in; F_ST_: Fixation index, measures the population differentiation due to genetic structure; LD: linkage disequilibrium; LD score: the sum of the LD r^2^ between the focal SNP and all the flanking SNPs within a 1cM window with 1000G data.

### Identifying and characterizing the function of cis-acting expression QTLs and splicing QTLs revealed common enrichment patterns between populations

We identified cis-eQTLs using QTLtools^34^(Fig. 1), adjusting for 20 hidden covariates identified by the probabilistic estimation of expression residuals (PEER)^35^ (Methods and Supplementary Note). These hidden factors were significantly correlated with technical and biological covariates such as experimental batch, RNA Integrity Number (RIN), sex, and age of death. We identified 3,278 genes with a cis-eQTL (false discovery rate (FDR) q-value < 0.05) in EAS populations, 10,043 genes with a cis-eQTL (FDR q-value < 0.05) in EUR populations, which are referred to as eGenes (Table 1).

**Table 1:** Identification of eQTLs and sQTLs

|  | #SNPs | #Genes | #Intron clusters | #cis-eQTLs/sQTLs | #Significant eGenes/sGenes |
| --- | --- | --- | --- | --- | --- |
| **EAS(n=145)** | 6,045,349 | 18,939 | 146,884 | 604,485/994,668 | 3,278/2,054 |
| **EUR(n=397)** | 8,108,028 | 16,542 | 188,310 | 2,790,193/4,705,755 | 10,043/5,641 |
| **Across population** | 4,681,303 | 16,266 | 132,619 | 286,288/442,281 | 2,650/1,779 |

Cis-eQTLs/sQTLs are defined as FDR q-value < 0.05. Significant cis-eQTLs/sQTLs across populations defined as eQTLs/sQTLs significant in EAS and EUR populations in the same direction. Significant eGenes are genes regulated by SNPs that passed multiple testing (FDR q-value < 0.05) based on the permutation-based analysis. Significant sGenes are genes that intron clusters can map to and passed multiple testing (FDR q-value < 0.05) based on the permutation-based analysis.

By identifying numerous excised intronic clusters using LeafCutter^36^ (Methods and Fig. 1), we were able to discover sQTLs as well. We identified 4,726 significant sQTLs (FDR q-value < 0.05) in EAS and 18,927 significant sQTLs (FDR q-value < 0.05) in EUR populations, which were mapped to 2,054 and 5,641 genes (sGenes) respectively per population (Table 1).

To better characterize the function of the eQTLs and sQTLs, we evaluated their distance distribution and enrichment in numerous functional regions. Our first finding agreed with previous results conducted in EUR brains^33,37,38^: 20% of the eQTLs in both populations were located within 10kb of the transcription start site (TSS) regions (Fig. 2a,b); the most significant (FDR_permutation_ q-value <0.05) SNP per sQTL (sSNP) showed clustering around the splice junction. Fifty percent of sQTLs are located within 10 kb of the splice junction (Fig. 3a,b) in both EAS and EUR populations, demonstrating that variants proximal to splicing junctions have a large effect. In contrast to eQTL, the majority of sSNPs (60%) lie within the gene body (Fig. 3c), also consistent with previous research^38^.

**Fig. 2.**
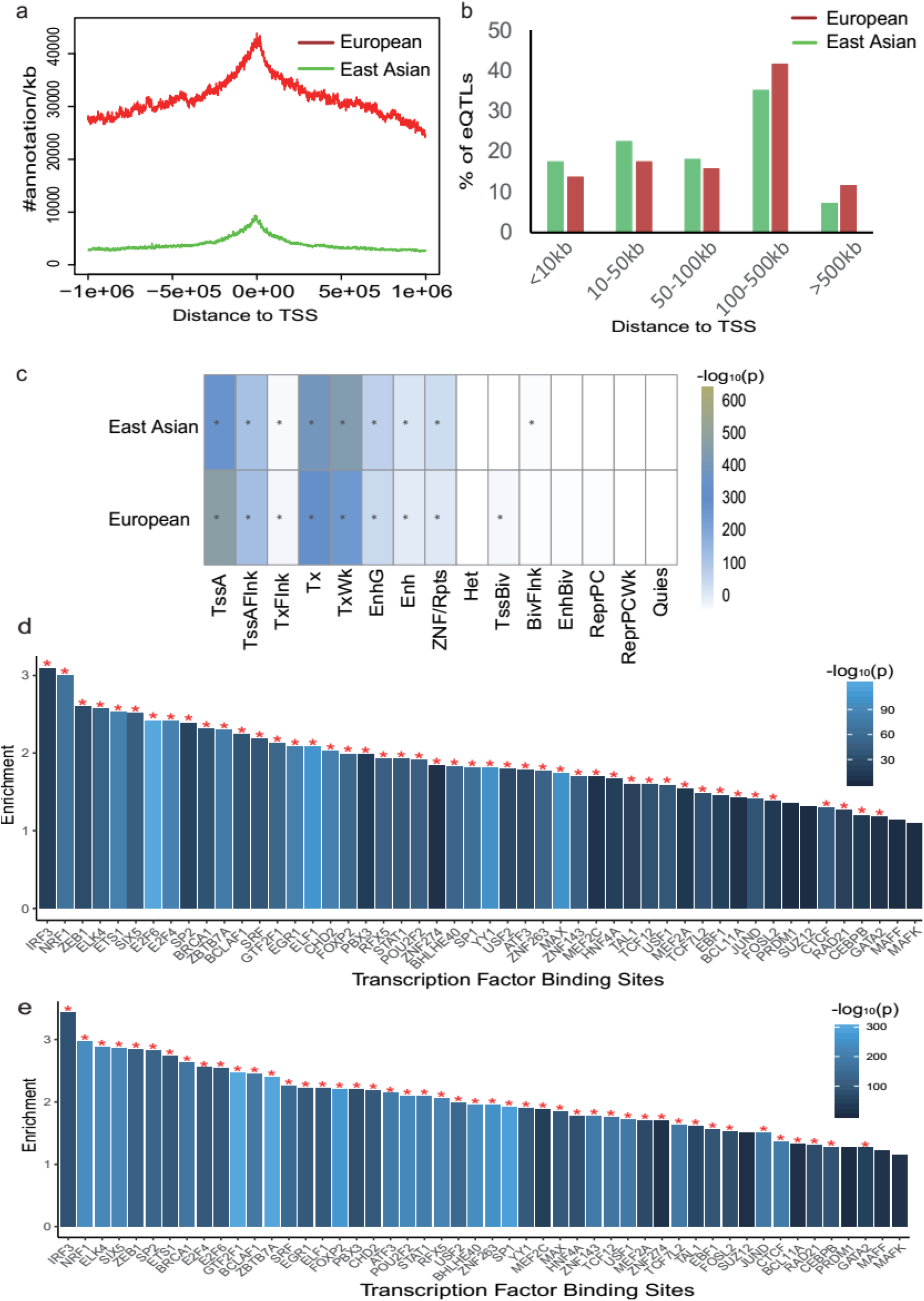
Characterization of eQTLs. **a,** Distance distribution of eQTLs in the East Asian (green) and European (red) populations to the TSS as defined in Gencode v19. **b,** Percentage of distance distribution of all cis-eQTLs in East Asian (green) and European (red) populations. **c,** Enrichment of eSNPs in 15 core models. eSNPs in both populations most significantly enriched in the TSSs, promoters, and transcribed regulatory promoters or enhancers. *P _Bonferroni_ < 0.05. **d,** Enrichments of eSNPs in experimentally discovered transcription factor binding sites in the East Asian population. **e,** Enrichments of eSNPs in experimentally discovered transcription factor binding sites in the European population. *P _Bonferroni_ < 0.05.

**Fig. 3.**
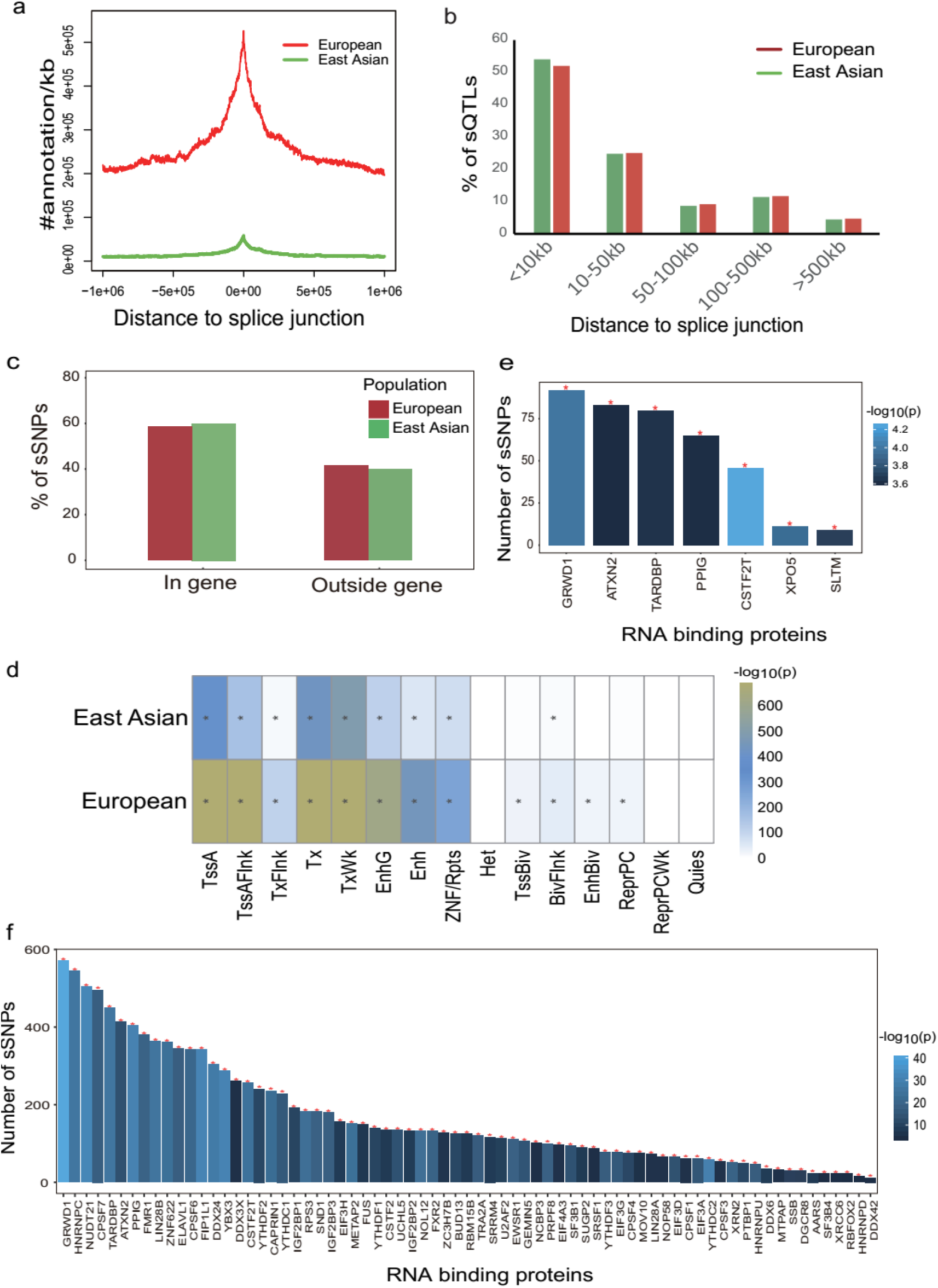
Characterization of sQTLs. **a,** Distance distribution of sQTLs to the splice junction. sQTLs from the East Asian (green) and European (red) populations. **b,** Percentage of distance distribution of all cis-sQTLs in East Asian (green) and European (red) populations. **c,** Fraction of sQTLs where the sSNP lies within vs outside its sGene. **d,** Enrichment of sSNPs in 15 core models. *P _Bonferroni_ < 0.05. **e,** RBP enrichment among the significant sSNPs in the East Asian population. **f,** RBP enrichment among the significant sSNPs in the European population. *P _Bonferroni_ < 0.05.

We then annotated expressed SNPs (eSNPs) and sSNPs with chromatin state predictions for prefrontal cortical tissue using GREGOR^39^ (Methods). We found that eSNPs and sSNPs were significantly enriched in the same TSSs, promoters, and transcribed regulatory promoters or enhancers (P_Bonferroni_ < 0.05, Fig. 2c, Fig. 3d; Supplementary Table 4). We also annotated eSNPs with transcription factor binding sites (TFBS) and sSNPs with experimentally determined RNA binding protein (RBP) binding sites. We observed that 46 and 49 TFBS were significantly enriched with eQTLs in the EAS and EUR populations separately (P_Bonferroni_ < 0.05, Fig. 2d,e and Supplementary Table 4). All of TFBS that were significantly enriched with eQTLs in EAS population were also significantly enriched with eQTLs in EUR population. Furthermore, sQTLs were significantly enriched in binding targets of 7 RBPs in the EAS population, while binding targets of 71 RBPs were significant in the EUR sQTL dataset (P_Bonferroni_ < 0.05, Fig. 3e,f and Supplementary Table 4). Five of the seven RBPs that were significantly enriched with sQTLs in the EAS population were also significantly enriched with sQTLs in EUR populations.

### Brain expression regulatory architectures are broadly preserved across EAS and EUR populations

An important aim in this study is to investigate to what degree the genetic control of gene expression and splicing in the brain varies between human ancestral populations. We first compared eGenes identified in EAS and EUR populations and found that 2,650 eGenes overlapped. Most (80%) eGenes detected in the EAS population were also significant in the EUR population (Fig. 4a). Additionally, we compared the eGenes detected in the EAS population to those from GTEx adult cortices^40^. We found 1,289 overlapping eGenes, accounting for nearly 40% between both datasets (Fig. 4b). We also compared sGenes across EAS and EUR populations. Results showed 1,779 overlapped sGenes (Fig. 4c), 87% of which were shared significantly across populations.

**Fig. 4.**
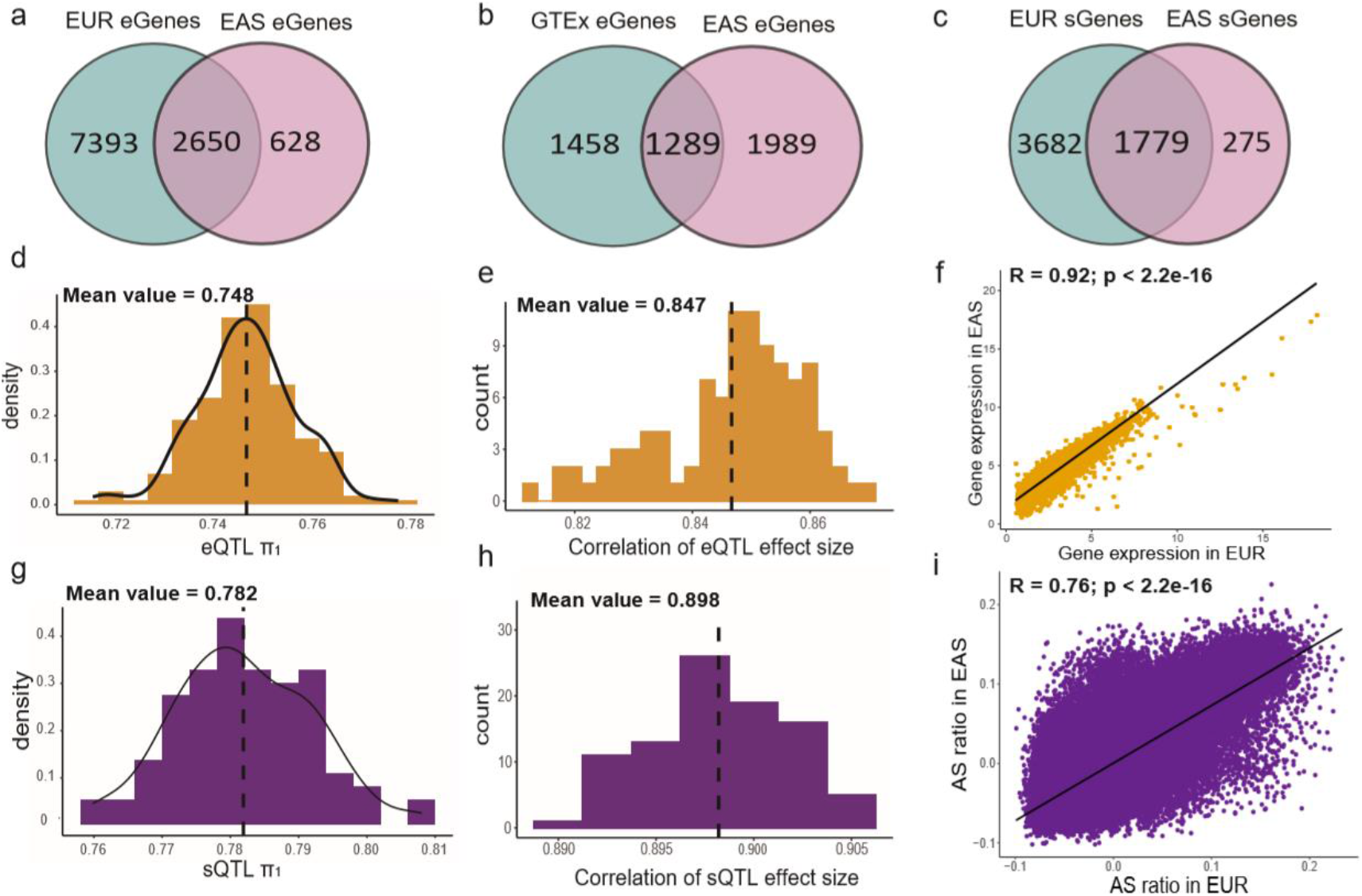
Comparison of the regulatory pattern. **a,** Venn Diagram for eGenes discovered in European (EUR) population vs East Asian (EAS) population. **b,** Venn Diagram for eGenes discovered in adult cortical tissue from GTEx vs EAS population. **c,** Venn Diagram for sGenes discovered in EUR population vs EAS population. **d,** Distribution of eQTL π_1_ between EAS and down-sampled EUR populations. The mean π_1_ value was 0.748. **e,** Distribution of correlation coefficient between eQTL effect size. The mean correlation coefficient value was 0.847. **f,** Pearson’s correlation in expressed genes between EAS and EUR populations. **g,** Distribution of sQTL π_1_ between EAS and down-sampled EUR populations. The mean π_1_ value was 0.782. **h,** Distribution of correlation coefficient between sQTL effect size. The mean correlation coefficient value is 0.898. **i,** Pearson’s correlation in intron clusters between EAS and EUR populations. AS: alternative splicing.

We next used Storey’s π_1_ statistic to assess the extent of eQTL/sQTL sharing across populations. To assess the true extent of this sharing, we performed down-sampling analysis with 100 repetitions (Methods). The fraction of eQTLs and sQTLs shared between the EAS and EUR populations was 74.8% and 78.2%, respectively (Fig. 4d,g). Moreover, we calculated the Pearson correlation of genetic effect size between shared QTLs and found that the genetic effect size between EAS and EUR populations was highly analogous (R_eQTL_=0.847; R_sQTL_=0.898; Fig. 4e,h).

We completed a meta-analysis, pooling results from diverse populations, to gain greater statistical power for QTL detection and to identify shared QTLs across populations. We calculated a meta p-value using METAL^41^ for each eQTL/sQTL across populations; eQTLs or sQTLs at a meta FDR < 0.05 were referred to as ‘meta eQTLs/sQTLs’. Greater than 80% of these were significant across populations and showed concordant regulatory direction across populations (Extended Data Fig. 2 and Supplementary Table 2). Also, numerous new eQTL/sQTL signals were identified by meta-analysis.

To comprehensively compare the brain expression regulatory architecture between EAS and EUR populations, we calculated the Pearson correlation of gene expression between the two populations using shared genes. We found that gene expression was highly correlated in the two populations (Fig. 4f; R = 0.92, p-value < 2.2e-16), and a similar result was observed for alternative splicing ratio (Fig. 4i; R = 0.76, p-value < 2.2e-16). Furthermore, we applied weighted gene co-expression network analysis (WGCNA)^42^ and robust WGCNA to create independent gene- and isoform-level networks. We then used preservation testing to evaluate the consensus of networks constructed by each population. Preservation Z summary score of each module was >2 in both the gene expression and isoform levels, showing that co-expression patterns are broadly preserved between EAS and EUR populations (Fig. 5c,d,e,f and Supplementary Table 5).

**Fig. 5.**
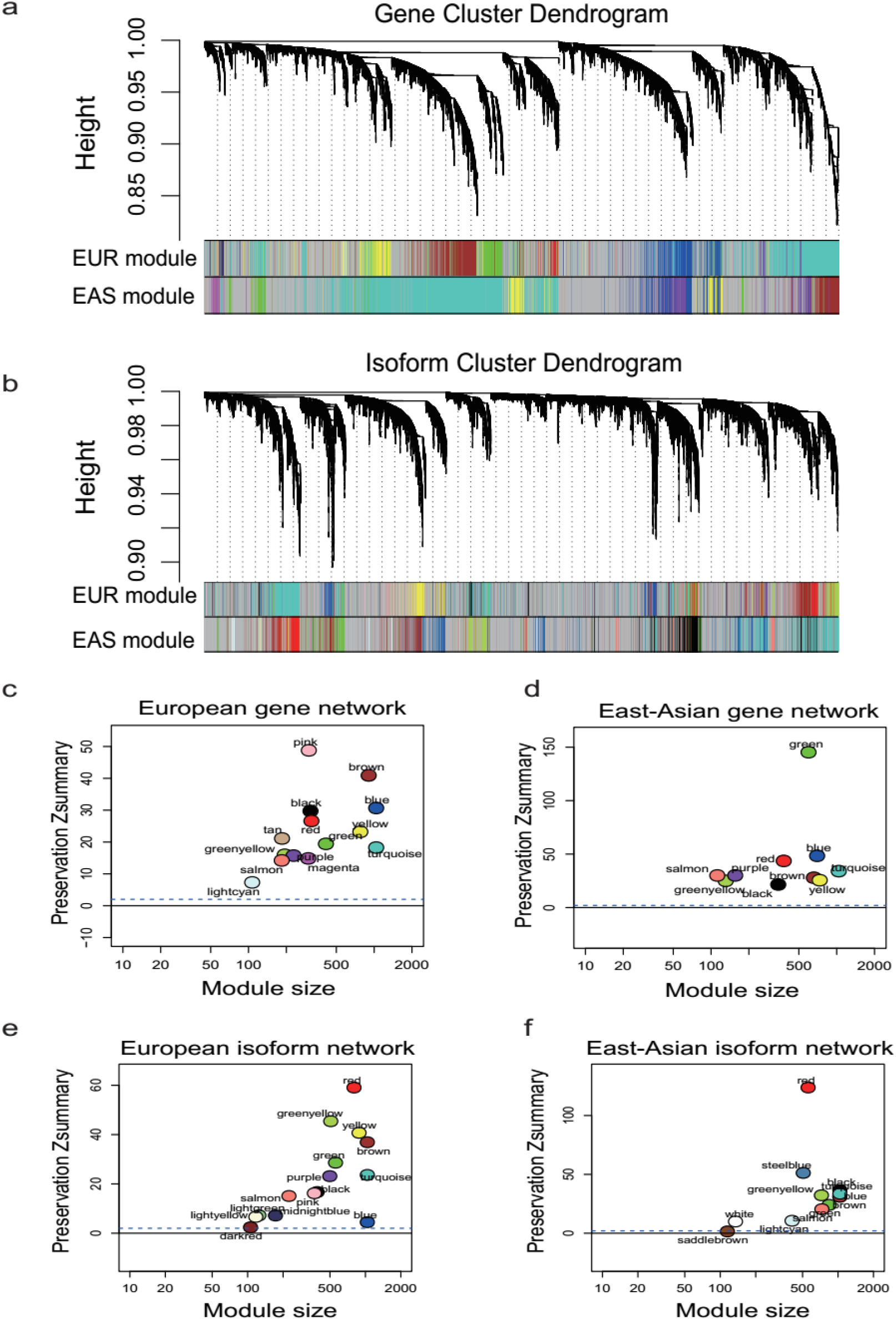
Comparison of co-expression pattern. **a,** Network analysis dendrogram based on hierarchical clustering of all genes by their topological overlap. Colored bars below the dendrogram show module membership. **b,** Network analysis dendrogram based on hierarchical clustering of all isoforms by their topological overlap. **c,** Preservation Z summary score of each gene co-expression module in the EUR population. The x-axis is the number of genes in each module and the y-axis is Z summary score, which measures the preservation between modules. When Z summary score >=2, it indicates that this module was preserved in another population. The blue dashed line is the moderately conserved threshold. Each point represents a module constructed in population, labeled by color. **d,** Preservation Z summary score of each gene co-expression module in the EAS population. **e,** Preservation Z summary score of each isoform co-expression module in the EUR population. **f,** Preservation Z summary score of each isoform co-expression module in the EAS population.

### Differences in AF and LD across populations decrease the QTL reproducibility and the power of gene expression prediction

Although the sharing ratio reached almost 80%, 20% of the QTLs are significant only in one population. We hypothesized that part of QTL differentiation is due to population divergence in AF and LD pattern across populations. To address this hypothesis, we defined EUR robust QTLs as significant QTLs detected at least 50 times in down-sampling analysis, ancestry-specific QTLs as significant in one population, and ancestry-shared QTLs as significant in both populations. We identified each of these QTL types by comparing the lists of EAS QTLs and EUR robust QTLs. To investigate the effect of AF on QTL differentiation, we estimated F_ST_ (fixation index) for each eSNP and sSNP and compared the distribution of F_ST_ between ancestry-specific and ancestry-shared QTLs (Methods). We found that ancestry-specific QTLs were significantly enriched in population-divergent SNPs (F_ST_ > 0.05; Fisher exact test: P < 2.2e-16) and ancestry-shared QTLs were significantly enriched in population-convergent SNPs (F_ST_ < 0.05; Fisher exact test: P < 2.2e-16; Fig. 6a). To further verify our hypothesis, we separated eSNPs/sSNPs into different minor allele frequency (MAF) bins, and calculated QTL π_1_ in each bin. We found that with similar AF, QTL sharing ratios were higher than cross SNPs with different AF, suggesting that the SNPs with less population divergence were more likely to be eQTLs/sQTLs shared by the two populations (Fig. 6b,c).

**Fig. 6.**
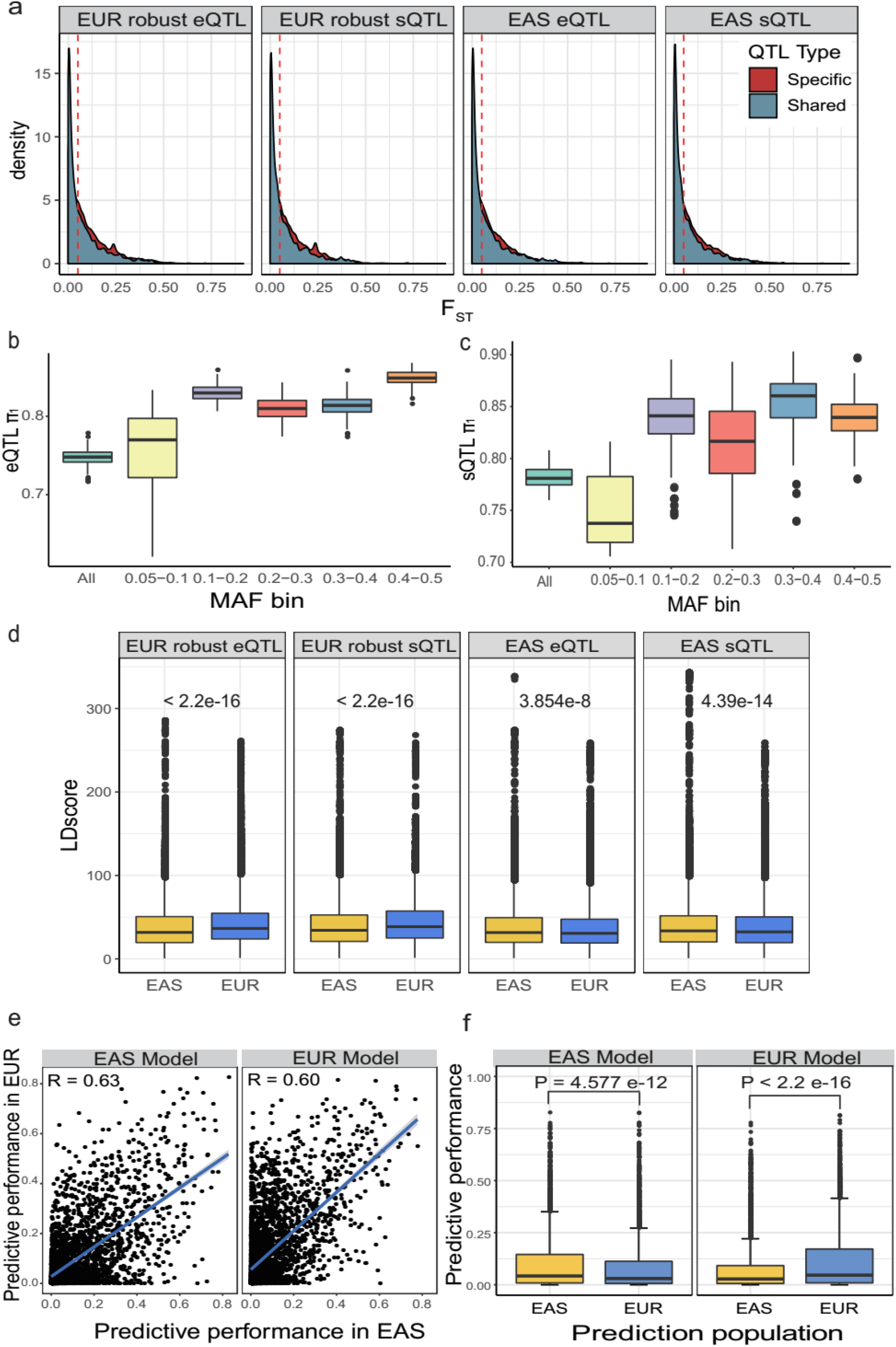
AF and LD differences contribute to QTL differences between populations. **a,** Comparison of F_ST_ between ancestry-shared and ancestry-specific QTLs. Robust QTL: detected as significant QTLs at least 50 times in down-sampling analysis. **b,** eQTL π_1_ in different MAF bins. **c,** sQTL π_1_ in different MAF bins. **d,** Linkage disequilibrium score distribution comparison for ancestry-specific QTLs between EAS and EUR populations. **e,** Comparison of predictive performance for each gene (R^2^) between EAS and EUR populations in different prediction models (EAS and EUR model). The identity line is shown in blue. **f,** Comparison of predictive performance between genes in different prediction models.

We then tested whether ancestry-specific QTL loci have unique LD patterns (Methods). Results showed LD patterns for ancestry-specific QTLs varied significantly between EAS and EUR populations (Wilcoxon tests, P < 0.05; Fig. 6d). Further, correlation coefficients between F_ST_ and LD score were less than 0.1 (P < 0.01), suggesting that the cross-population differences in LD patterns affect QTL differentiation independently when compared with F_ST_.

Associations between SNPs and genes enable the development of predictive models that can “impute” gene expression when phenotype-related tissue types are unavailable. However, population-specific QTL signals may reduce the accuracy of gene expression prediction across populations. We hypothesize that prediction performance will be lower when a model trained on one population is used to predict gene expression in another population. To investigate, we compared gene-expression predictive performance within and across EAS and EUR populations. We used matched SNPs and genes in both populations (n=145) to build the models (n = 100) using PrediXcan^15^ (Methods). The Pearson correlation between predictive performance of genes in the EAS and EUR populations was 0.60 (Fig. 6e). We also found that single-population-trained models had significantly decreased performance when predicting gene expression in another population (Wilcoxon tests, EAS Model: P = 4.577 × 10^-12^; EUR Model: P < 2.2 × 10^-16^; Fig. 6f; Supplementary Table 3)

### Synapse- and GTPase-related pathway implicated in SCZ risk across populations

eQTLs and sQTLs detected in the human brain can help to decipher the unlock biological mechanisms of SCZ. To examine whether QTL results from the phenotype-linked population explains more disease signals and SNP heritability than those from disparate populations, we first collected SCZ GWAS summary statistics for both ancestral populations from the Psychiatric Genomics Consortium (PGC)^43,44^. We then compared GWAS signal enrichment using partitioned LD-score regression (LDSR)^45^ (Methods). For the EAS GWAS summary data, eQTLs/sQTLs detected in the EAS population showed greater significant enrichment in GWAS signals than those from the EUR population (Welch Modified Two-Sample t-Test P < 0.001) and vice versa (Welch Modified Two-Sample t-Test P < 0.001; Fig. 7a and Supplementary Table 6). We corrected for possible sample size variance bias by calculating the enrichment of robust EUR QTLs, and the results agreed with previous reports (Extended Data Fig. 3a,b).

**Fig. 7.**
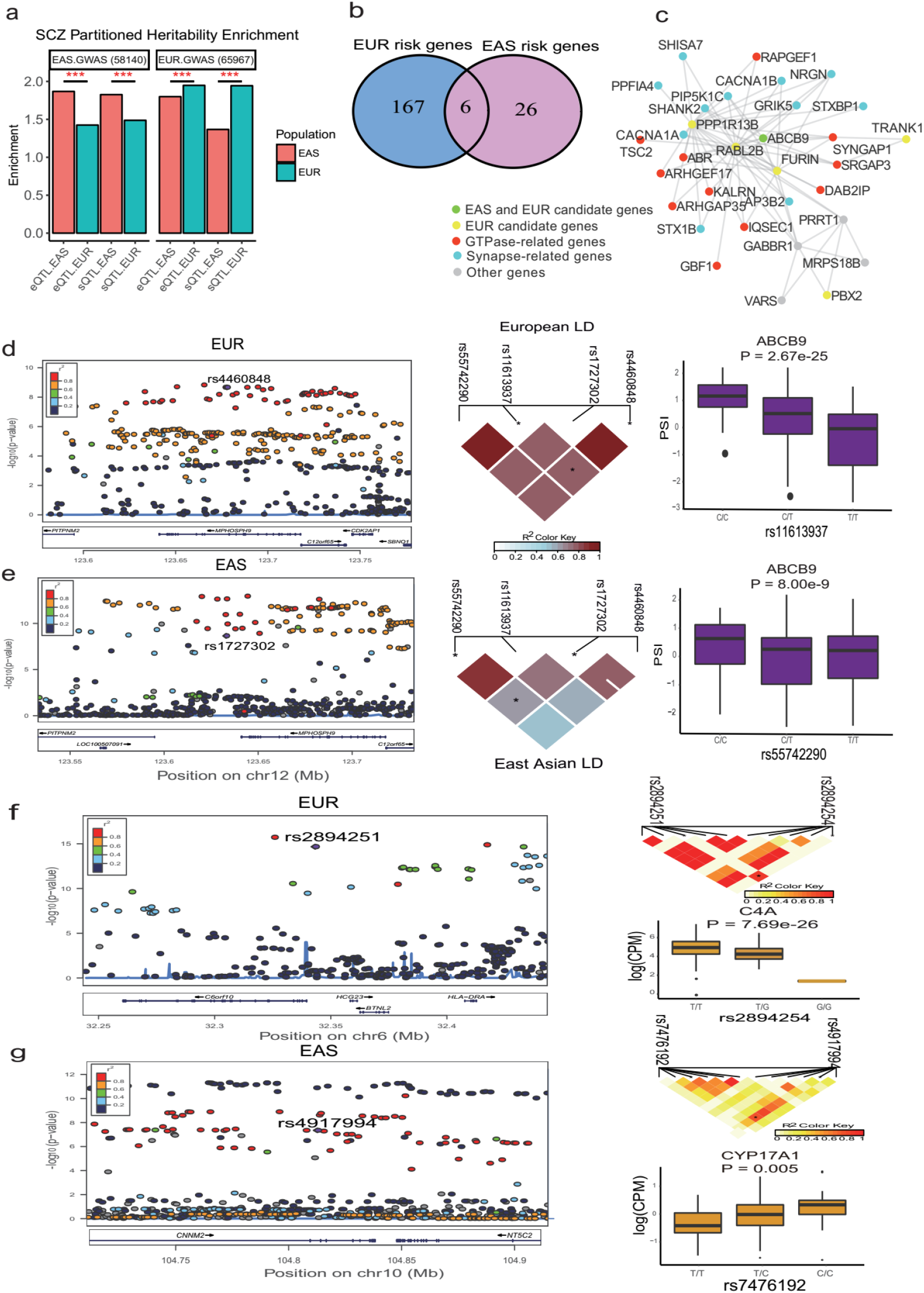
Explanation of SCZ GWAS signals and prioritization of candidate genes. **a,** GWAS enrichment results from LDSR. ***: Welch Modified Two-Sample t-Test P < 0.001. **b,** Venn plot for SCZ risk genes in EUR and EAS populations with combined RTC and SMR results. **c,** The sub-network of yellow module. **d-e,** Examples of shared SCZ risk genes. **f,** Example for EUR-specific risk gene, C4A. **g,** Example for EAS-specific risk gene, CYP17A1. LDSR: LD score regression.

It was next necessary to evaluate whether observed differences represented true etiologic heterogeneity of SCZ across populations. To achieve this, we used regulatory trait concordance (RTC)^46^ and summary data-based Mendelian randomization (SMR)^9^ to prioritize SCZ candidate risk genes (Methods). We prioritized 199 SCZ candidate risk genes, including 173 genes in the EUR population and 32 in the EAS population (Supplementary Table 7; Supplementary Table 8 and Fig. 7b). Six of the 199 were identified in both populations (CNNM2, C12orf65, MPHOSPH9, MARCKSL1P1, C2orf47, and ABCB9). Other genes were identified within a single population. For example, C4A was identified as a risk gene in the EUR population (Fig. 7f) by integrating the eQTL signals, while CYP17A1 was identified as a risk gene in the EAS population by integrating the eQTL signals (Fig. 7g). Comparing published SCZ risk genes with the 199 candidate genes we identified, 77 of the EUR candidate genes and 10 of the EAS candidate genes aligned (Methods and Supplementary Table 9).

Along with peripheral genes, these candidate genes form a network that fulfills specific functional roles. To better characterize the biological function of these candidate genes, we analyzed the enrichment of candidate genes having previously constructed networks (Methods). We tested whether modules were significantly enriched in candidate genes for both EAS and EUR populations, but none were. We also tested whether any consensus modules (preserved in both populations) were significantly enriched with candidate genes present in both populations. One consensus module was significantly enriched in candidate genes from the combined populations (Fig. 7c; p-value = 0.01). Enrichment analysis showed function related to synapse and GTPase pathways, including regulation of chemical synaptic transmission, neuron projection development, synapse structure or activity, small GTPase mediated signal transduction, and GTPase binding (Extended Fig. 3d and Supplementary Table 10).

## Discussion

We developed a novel brain transcriptome data set and compiled the first genome-wide brain regulatory map of the prefrontal cortex from a solely EAS population. We identified 3,278 eQTL and 4,726 sQTL signals that reached a genome-wide level of significance. Detected eSNPs and sSNPs corresponded to previous reports^38,40,47,48^ with significant enrichment in active functional regions such as promoters and enhancers. Comparing the EAS data with the PsychENCODE/BrainGVEX-derived EUR data, we found most regulatory elements common to both populations. Moreover, by integrating QTL signals with summary statistics from SCZ GWAS, we observed synapse- and GTPase-related pathways involved in the development of SCZ in both populations.

This study demonstrated convergent transcriptional regulatory architectures between EAS and EUR populations through multiple lines of evidence. Meta-analysis revealed approximately 80% of QTLs were shared between populations. Moreover, several relational analyses suggest a high degree of congruence between EAS and EUR populations, including the π_1_ statistic (eQTL=0.748; sQTL=0.782), correlations between populations for genetic effect size (eQTL=0.847; sQTL=0.898) and correlations for gene expression and co-regulatory networks. Our study of post-mortem brain tissue concurs with studies based on whole blood and liver tissue in which EAS and EUR cis-eQTL replication rates equaled 60%^49^ and 40%^50^ respectively. Our study parallels previous comparisons of the genetic control of gene expression^24,50,51^, methylation^52^, and chromatin accessibility^53^ generated from lymphoblastoid cell lines in diverse populations from worldwide reference panels^54–56^, showing that regulatory patterns are shared across populations.

Our proposal that some QTLs have population-specific effects is not unique ^24,53^. Seeking to find the biological mechanisms underlying these divergent effects, we compared F_ST_ and LD score distribution for ancestry-specific QTLs across populations. Here, we found significant differences. These results indicate that a degree of QTL differentiation signals divergence in AF and LD. Evidence suggests that such genetic differentiation in ancestral populations is due primarily to natural selection^57,58^. Besides, contemporary populations descend from dramatically smaller migratory populations (bottleneck effect), hence, population-specific QTLs could arise from bottleneck effect and environmental factors including climate, diet, and pathogenic microorganisms.

Population-specific QTLs have important implications for predictive modeling. QTL signals lay the foundation for predictive models and assist in imputing gene expression when tissues relevant to phenotypes are unavailable. Our gene prediction model that was trained in one population decreased their prediction performance when predicting gene expression for other populations at a ratio of 14% to 33% respectively. This agrees with several recent studies reporting superior accuracy of prediction models in target populations with ancestry comparable to the discovery population^14,21–23^. Therefore, population-specific predictive models are integral for transcriptome mapping of the human brain.

Population-specific regulatory regions may harbor a portion of the disease risk. This may limit QTL’s utility in interpreting GWAS signals in disparate populations. We compared the enrichment of QTL signals in SCZ GWAS across populations and found more significant enrichment of eQTLs/sQTLs in the discovery population than in the disparate populations. Similar results have been reported in Type 2 diabetes^59^. These findings highlight the importance of using GWAS to interpret QTLs from the target population in accurately explaining the disease signals within that population.

Since SCZ occurs with similar prevalence and a genetic basis broadly shared across populations^43,60^, we would expect distinct groups to share many risk genes. Surprisingly, we observed only six of the 199 SCZ risk genes in common between EAS and EUR populations. Nevertheless, this finding should not be interpreted to mean that EAS and EUR populations carry entirely different risk genes. For, we found that almost 70% of the SNPs that are linked to regulating population-specific SCZ risk genes vary in AF and LD patterns across populations. For example, we identified the EUR-specific risk gene C4A, a target of extensive scrutiny in association with SCZ^61–63^. C4A localizes to the MHC class III region on chromosome 6, which is strongly associated with SCZ and which hosts a EUR-specific LD pattern. The corresponding GWAS signal rs2894251 was significant within the EUR population (P = 2.144 e-15; MAF = 0.12) but not so in the EAS population (P = 0.05156; MAF = 0.02). Although the associated eSNP rs2894254 is extremely uncommon in the EAS population (MAF < 0.001), it has an MAF of 6% in the EUR population. These results suggest that at least some differences in EAS and EUR SCZ risk genes are due to low AF and disparate LD patterns, which may account for the loss of risk genes. Risk genes were more readily detectable in both populations when they were present in similar or higher AFs with similar LD patterns. Yet, many SCZ risk genes are evident within low frequencies too, hindering consistent detection.

It is well known that complex molecular networks and cellular pathways fuel disease susceptibility and development^64,65^. Therefore, we exploited pathway enrichment analyses of identified risk genes and co-regulated genes to explore the mechanisms behind SCZ. SCZ risk genes were significantly enriched in one consensus module (module yellow) for both ancestral populations. This module was enriched for an array of established SCZ modular pathways, including synapse- and GTPase-related pathways^66–70^. Pathways related to neuron-to-neuron, postsynaptic density, and asymmetric synapses are established suspects in genetic risk of SCZ. A meta-analysis showed a significant decrease in the density of postsynaptic elements in SCZ patients compared to healthy controls^69^. GTPase-related pathways, including regulation of small GTPase-mediated signal transductions and GTPase binding, have also been implicated in SCZ. A previous study shows that a missense polymorphism (H204R) of a Rho GTPase-activating protein is associated with schizophrenia in men^70^. Our results support the premise that synapse-related and GTPase-related pathways have an important role in the etiology of schizophrenia for both EAS and EUR populations.

Our study showed how both eQTLs and sQTLs benefit the study of underlying disease mechanisms. We discovered how heritability explained by eQTLs and sQTLs is similar in both EAS and EUR populations. Recent studies have examined the contribution of regulatory variants to SCZ, educational attainment, and autism spectrum disorder (ASD), concluding that sQTLs contribute comparably or with even greater magnitude than eQTLs^8,14,38^. Additionally, we found that although only 14% of SCZ risk genes were identified by both eQTL and sQTL signals (Extended Data Fig. 3c,d), 40% of the SCZ risk genes identified by integrating either eQTLs or sQTLs had also been reported as SCZ risk genes in previous literature^66,71–77^. This result indicates that eQTLs and sQTLs can identify distinct risk genes which facilitate our understanding of disease mechanisms.

In general, our results show the transcriptional architecture of expression regulation and the underlying SCZ biology converging between the EAS and EUR populations. Synaptic- and GTPase- related pathways are likely suspects in the pathogenesis of SCZ in both populations. Future studies should assemble a range of large samples from worldwide ancestral populations to establish whether these findings are applicable globally. If so, mechanistic studies could narrow in on fewer pathways toward extracting the pathogenesis of SCZ with greater precision.

## Methods

### EAS sample collection, sequencing and EUR public data collection

We collected 151 prefrontal cortical samples of Han Chinese descent from the National Human Brain Bank for Development and Function according to the standardized operational protocol of China Human Brain Banking Consortium^78,79^, and under the approval by the Institutional Review Board of the Institute of Basic Medical Sciences, Chinese Academy of Medical Sciences, Beijing, China (Approval Number: 009-2014). We sequenced 151 samples following the BGISEQ-500 protocol outsourced to BGI, WGS and transcriptome sequencing was performed on BGISEQ-500 platform with an average depth of 10X (Supplementary Table 1 and Supplementary Note). To assess differences in ancestry, we also downloaded and processed raw whole genome and RNA-seq data for 407 European ancestry from PsychENCODE/BrainGVEX (Synapse number: syn4590909).

### EAS data quality control

Raw sequencing reads were filtered to get clean reads by using SOAPnuke (v1.5.6)^80^, and used FastQC to evaluate the quality of sequencing data via several measures, including sequence quality per base, sequence duplication levels, and quality score distribution for each sample. The average quality score for overall DNA and RNA sequences show high scores above 30, indicating that a high percentage of the sequences had high quality (Supplementary Note).

### Variant identification

Clean DNA sequencing reads were mapped to the human reference genome hg19 (GRCh37) using BWA-MEM algorithm (BWA v. 0.7.128)^81^. Ambiguously mapped reads (MAPQ <10) and duplicated reads were removed using SAMtools v. 1.29^82^ and PicardTools v. 1.130 (http://picard.sourceforge.net/) respectively. Genomic variants were called following the Genome Analysis Toolkit software (GATK v. 3.4–46) best practices^83^. The ancestry of each sample was inferred using data from the 1000 Genomes Project, and no sample was excluded. For EAS cohort, genotypes were imputed into the 1000 Genomes Project phase 3 EAS reference panel by chromosome using Michigan Imputation Server^84^ and subsequently merged. Imputed genotypes were filtered for R^2^ < 0.3, Hardy-Weinberg equilibrium p-value < 1 × 10^-6^ and MAF < 0.05, resulting in ~6.4 million autosomal SNPs. For EUR cohort, genotypes were imputed into the HRC reference panel, and removed SNPs with R^2^< 0.3, HWE p-value < 1 × 10^-6^ or MAF < 0.01.

### Gene-expression quantification and filter

Mapping of RNA-sequencing reads was completed using STAR (2.4.2a)^85^ and the quantification of genes and transcripts was with RSEM (1.3.0)^86^. Raw read counts were log-transformed by R package VOOM first^87^, filtering those with log2(CPM)<0 in more than 75% of the samples. We removed all transcripts derived from mitochondrial DNA and X and Y chromosomes. Samples with a Z-score (assessing connectivity between samples) lower than −3 were removed. Quantile normalization was then used to equalize distributions across samples.

### Intron cluster quantifications

We used Leafcutter to quantify clusters of variably spliced introns^36^. A cluster consists of overlapping introns that share a splice site. The usage of each intron was first quantified using previously aligned FASTQ files from STAR. Overlapping introns were then grouped with the settings of 50 reads per cluster and a maximum intron length of 500kb.

### Co-expression network analysis

To place results from individual genes within their systems-level network architecture, we performed WGCNA^42^ using human brain RNA-seq data. Individual (covariate-regressed) expression datasets were combined using the 16,266 genes present across all studies. The resulting normalized mega-analysis expression set was used for all downstream network analyses. We also using robust WGCNA (rWGCNA) to reduce the influence of potential outlier samples on the network architecture. Module robustness was ensured by randomly resampling (2/3 of the total) from the initial set of samples 100 times. This was followed by consensus network analysis, a meta-analytic approach to define modules using a consensus quantile threshold of 0.2. The parameter of rWGCNA was consistent with normal WGCNA (Supplementary Note).

### eQTL and sQTL mapping

We used PEER^35^ to identify hidden confounders and evaluated the correlation between the known factors (such sex and age) with hidden confounders. We then performed cis-eQTL and cis-sQTL mapping using QTLtools^34^, adjusting for PEER factors (Supplementary Note), with a defined cis window of one megabase up- and downstream of the gene/intron cluster body. QTLtools was run in nominal pass mode to detect all available QTLs. QTLtools was also run in the permutation pass mode to identify the best nominal associated SNP per phenotype and with a beta approximation to model the permutation outcome. P-values were then multiple testing corrections using the “q-value” package in R. We define FDR q-value < 0.05 as significant QTL.

### Functional enrichment

We performed functional enrichment of both eQTLs and sQTLs using GREGOR^39^ (Genomic Regulatory Elements and Gwas Overlap algoRithm) to evaluate the enrichment of variants in genome-wide annotations. GREGOR calculated the enrichment value based on the observed and expected overlap within each annotation. We downloaded the 15-state ChromHMM model BED (Browser Extensible Data) files from the Roadmap Epigenetics Project^88^. We also downloaded 78 consensus transcription factor and DNA-protein binding site BED files existing in multiple cells^89^ and then filtered to 50 binding proteins that showed cortical brain expression in EAS and EUR populations data. Lastly, we obtained 171 human RBP site BED files from POSTAR2 database, which was developed as the updated version of CLIPdb and POSTAR and provides the largest collection of RBP binding sites and functional annotations^90^.

### The fraction of shared eQTL/sQTL between EAS and EUR population

Sharing rate was assessed form all significant eQTLs/sQTLs in the discovery dataset by estimating the proportion of true associations (π_1_) on the distribution of corresponding p-values of the overlapping eQTLs /sQTLs in the replication dataset^93^. To avoid the influence of sample size in pairwise comparison and get the true replication rate, we randomly selected a subset of European samples (n=151), followed the same pipeline to detect QTLs, and calculated the correlation of genetic effect size of shared eQTL/sQTL between EAS and EUR populations, repeating 100 times. We calculated π_1_ and used the mean value of π_1_ to assess reproducibility between EAS and EUR.

### Network preservation analysis

To generate population-specific networks, we compared networks between constructed EAS and EUR populations by individual. We then used WGCNA-integrated function (modulePreservation) to calculate module preservation statistics and applied the Z summary score (Z-score) to evaluate whether a module was conserved or not.

### F_ST_ analysis

We used the EAS and EUR panels from the 1000 Genomes Project Phase 3 to investigate the Fixation index (F_ST_). We estimated F_ST_ using vcftools^91^ following the Weir and Cockerham approach^92^ for each eSNP and sSNP.

We defined population-divergent SNPs as those with F_ST_ >= 0.05 and population-shared SNPs as those with F_ST_ < 0.05. To collect the list of ancestry-specific QTLs and ancestry-shared QTLs, first, we defined EUR robust QTLs as those that were called significant at least 50 times in down-sampling analysis, as well as ancestry-specific QTLs (significant in one population) and ancestry-shared QTLs (significant in these two populations), by comparing the list of EAS QTLs and EUR robust QTLs. Finally, we performed Fisher’s exact test between ancestry-specific QTLs and population-divergent SNPs, as well as population-shared SNPs and ancestry-shared QTLs to test the contribution of AF in QTL comparison.

### LD pattern comparison

We calculated the LD score for each SNP as the sum of the LD r^2^ between the focal SNP and all flanking SNPs within a 1cM window within the corresponding 1000G EAS and EUR genotype data. We then mapped ancestry-specific eSNPs and sSNPs into LD-score files to obtain the LD score for each ancestry-specific eSNPs or sSNPs in each population. We then performed Wilcox testing to evaluate whether the mean value of the LD score was significantly varied between populations.

### Gene-expression prediction

We used matched SNPs and genes from the EAS and EUR populations using matching sample sizes (n=145) to build the gene-expression prediction model. We separated each population into training and validation datasets (100 for training and 45 for validation). Prediction models were built using PrediXcan^15^ (Elastic Net) for both populations. Predictive performance (R^2^) was measured within each population using nested cross-validation. Wilcoxon tests measured any significant difference in prediction performance across populations.

### Partitioned LDSC

Partitioned LD score regression v1.0.1 was used to measure the enrichment of GWAS summary statistics in each functional category by accounting for LD^45^. Brain QTL annotations were created by eSNP and sSNP, mapped to the corresponding 1000 Genome reference panel. LD scores were calculated for each SNPs in the QTL annotation using an LD window of 1cM in 1000 Genomes European Phase 3 and 1000 Genomes Asian Phase 3 separately. Enrichment for each annotation was calculated by the proportion of heritability explained by each annotation divided by the proportion of SNPs in the genome falling in that annotation category. We then applied Welch Modified Two-Sample t-Test on enrichment values generated from QTLs in the two populations.

### Colocalization

We used the conditional association as described in Nica *et al.*^46^ to test for evidence of colocalization. This method compares the p-value of association for the lead SNP of an eQTL or sQTL before and after conditioning on the GWAS hits. The equation for the regulatory trait concordance (RTC) Score is as follows: RTC= (N_SNPs_ in an LD block/Rank_GWAS_SNP_)/ N_SNPs_ in an LD block. The rank denoted the number of SNPs, which when used to correct the expression data, have a higher impact on the QTL than the GWAS SNPs (i.e., Rank_GWAS___SNP_ = 0 if the GWAS SNP is the same as the eQTL or sQTL SNP and Rank_GWAS_SNP_ = 1 if, of all the SNPs in the interval, the GWAS SNP has the largest impact on the eQTL or sQTL). RTC values close to 1.0 indicated causal regulatory effects. A threshold of 0.9 was used to select causal regulatory elements.

### Prioritizing genes underlying GWAS hits

We applied an SMR^9^ method on EAS and EUR SCZ GWAS summary data to prioritize candidate genes. We used nominally significant QTLs identified in the previous analysis (FDR < 0.05), containing thousands of unique probes with filtered GWAS summary data (p < 0.01) to perform the SMR test. In general, we use the default parameters suggested by the developers of the SMR software. These included the application of heterogeneity independent instruments (HEIDI) testing, filtering out hits that arose from significant linkage with pleiotropically associated variants (LD cutoff of P = 0.05 in the HEIDI test, as suggested by SMR). Genes with an empirical P passed Bonferroni correction in the SMR test and a P > 0.05 in the HEIDI test were considered as risk genes.

### Schizophrenia-related signals

The schizophrenia risk gene sets were collected from publications and databases. For gene analysis, we collected these genes and converted them to Ensembl Gene IDs in Gencode (hg19) using BioMart. We examined whether the risk genes meet one of these criteria: (1) affected by copy number variants (CNVs)^71^; (2) identified by linkage and association study^72–74^; (3) had de novo variants from NP de novo database^75^; (4) identified by convergent functional genomics (CFG)^76^; (5) identified by Pascal gene-based test^76^; or (6) expressed differentially in SCZ^66,77^.

### Module enrichment

Module functional enrichment of Gene Ontology pathways was assessed with GO-Elite v1.2.5^93^ as well as using the clusterprofiler^94^ R package, using GO and KEGG databases. For gProfiler, “moderate” hierarchical filtering was used. A custom background set consisted of 10,387 genes present across all studies and microarray platforms. The top pathways were those reaching significance with FDR-adjusted P < 0.05. Module eQTL and candidate genes enrichment were assessed with Fisher exact testing in R.

## Acknowledgments

We thank all donors and their families. We thank professor Hailiang Huang for sharing the EAS SCZ GWAS summary statistics. We thank Dr. Elliot Gershon for helpful comments. Tissue was provided by the Human Brain Bank, Chinese Academy of Medical Sciences & Peking Union Medical College, Beijing, China, the Chinese Brain Bank Center, and the Xiangya School of Medicine Brain Bank. This study was supported by National Natural Science Foundation of China (Grants Nos. 31571312, 31970572, 31871276, 91632116 and 81401114), the National Key R&D Project of China (Grants No. 2016YFC1306000 and 2017YFC0908701), Innovation-driven Project of Central South University (Grant Nos. 2020CX003, 2015CXS034 and 2018CX033), Hunan Provincial Natural Science Foundation of China (Grant No. 2019JJ40404), the Fundamental Research Funds for the Central Universities of Central South University, and NIH grants 1 U01 MH103340-01, 1R01ES024988.

## Author contributions

S.L. drafted the manuscript, performed the genotype and RNA-seq quality control, mapped eQTL, performed functional enrichment, compared the brain regulatory architecture, as well as integrated QTLs with GWAS signals. Y.C. wrote the manuscript, constructed co-regulatory networks, performed preservation test as well as pathway enrichment analysis. F.W. wrote the manuscript, performed sQTL mapping and functional enrichment analysis. Y. J. preprocessing the genotype and RNA-seq data from PsychENCODE/BrainGVEX project. F.D. and M.L. extracted DNA and RNA, as well as collected sample information. Y.X., R.K, and L.K. substantively revised the manuscript. Z.N. and S.X. participated in the design of comparing the brain regulatory architecture. Sample provided by W.Q., C.M., X.Y., A.B., J.D., J.H. and B.T. C.L. provided PsychENCODE/BrainGVEX data. C.C. conceived, designed and supervised the study and modified the manuscript.

## Data availability

The raw sequence data reported in this paper have been deposited in the Genome Sequence Archive in BIG Data Center, Beijing Institute of Genomics (BIG), Chinese Academy of Sciences, under accession numbers HRA000108, HRA000108 that can be accessed at https://bigd.big.ac.cn/gsa-human. eQTL and sQTL summary results for EAS samples can be downloaded from http://brainexpnpd.org:8088/BrainEXPNPD/download.html.

## Code availability

Codes are available at https://github.com/liusihan/population-compare-pipeline

## Additional information

Correspondence and requests for materials should be addressed to C.C.

**Extended Data Fig.1.**
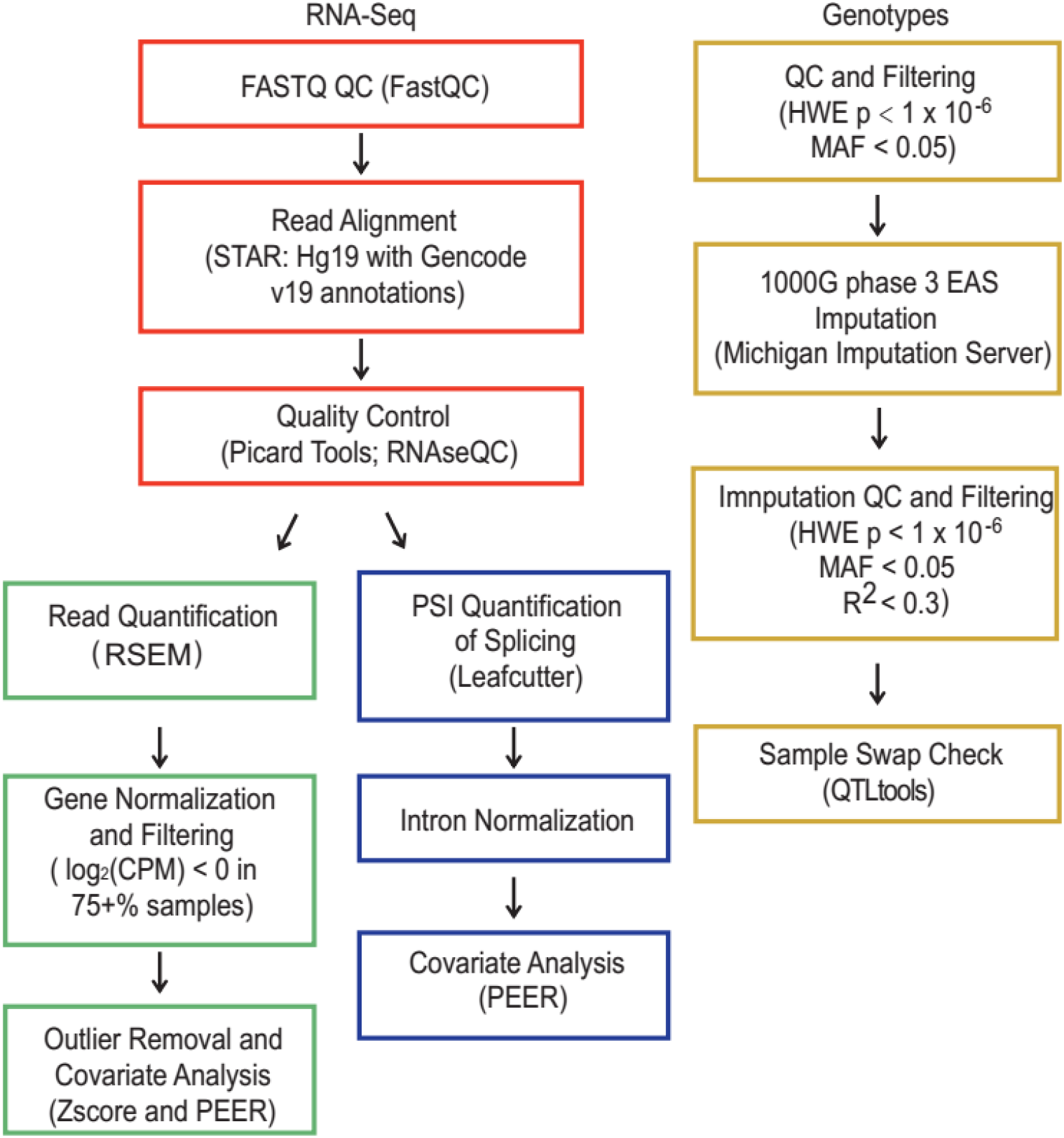
Overview of methods and QC pipeline for EAS samples.

**Extended Data Fig.2.**
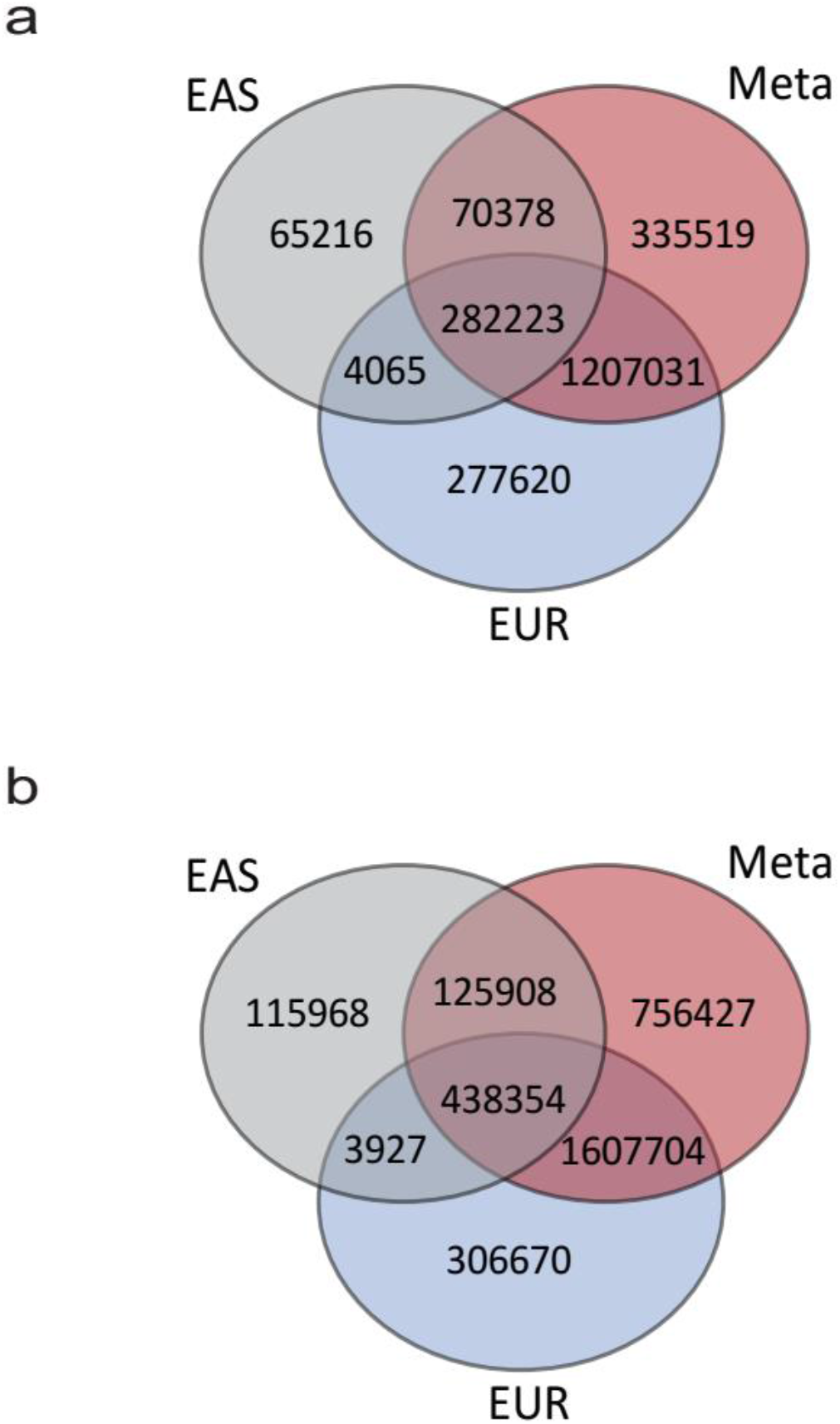
QTL comparison. **a,** Venn plot for eQTLs. **b,** Veen plot for sQTLs.

**Extended Data Fig.3.**
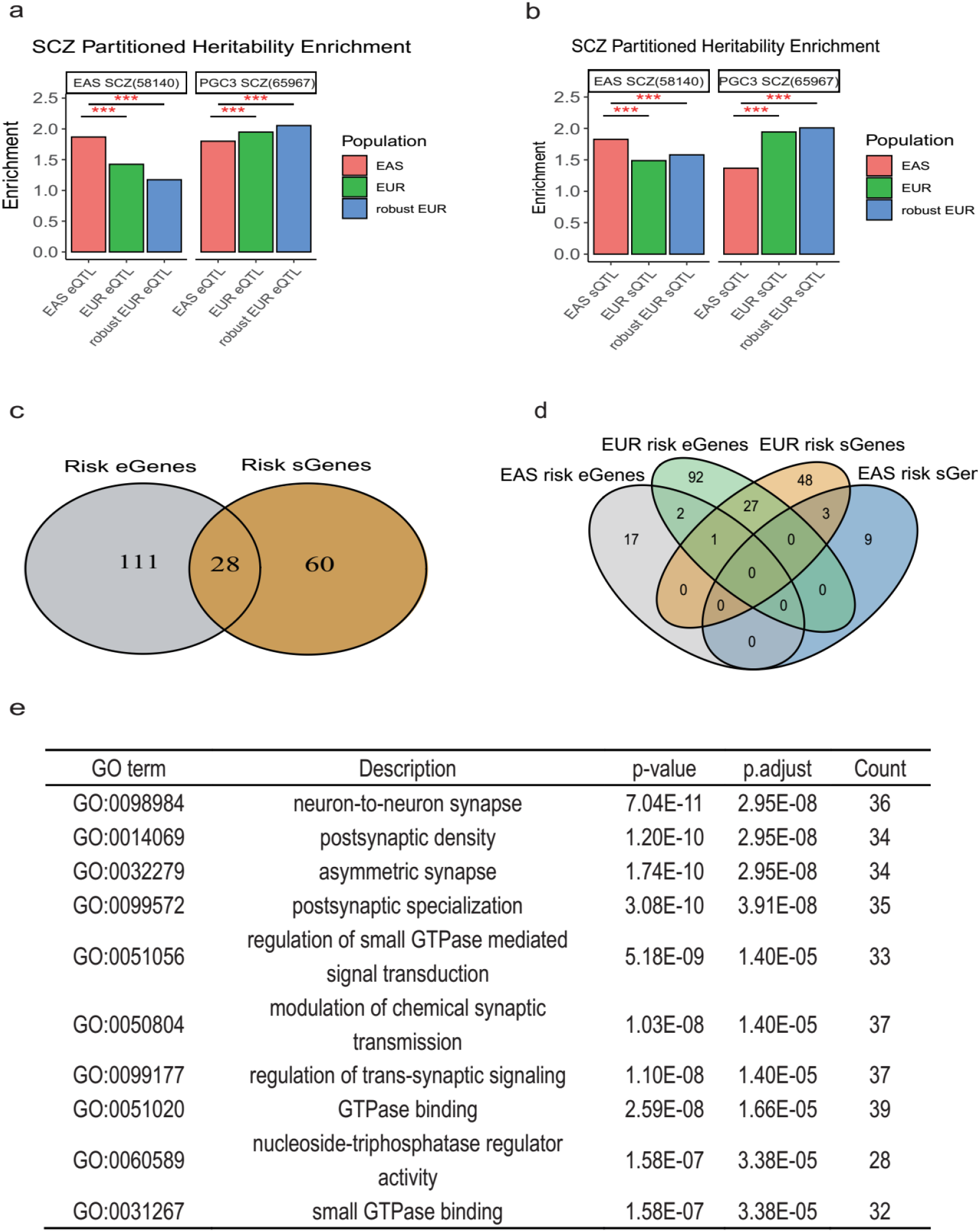
Integrating SCZ GWAS signals. **a,** GWAS signals enrichment comparison for eQTLs. **b,** GWAS signals enrichment comparison for sQTLs. **c-d,** Venn plot for risk genes identified by eQTLs and sQTLs. **e**, List of the top ten pathways which enriched in SCZ risk genes. p.adjust: Bonferroni adjusted p-value. Count: number of genes located in this pathway.

